# Urban *Cepaea nemoralis* snails are less likely to have nematodes trapped within their shells

**DOI:** 10.1101/2024.03.07.583959

**Authors:** Maxime Dahirel, Hannah Reyné, Katrien De Wolf, Dries Bonte

## Abstract

Urbanization is a major human-induced environmental change which can impact not only individual species, but also the way these species interact with each other. As a group, terrestrial molluscs interact frequently with a wide diversity of parasites, yet the way these interactions vary across space and in response to environmental pressures is poorly documented. In this study we leveraged a recently discovered defence mechanism, by which snails trap parasitic nematodes in their shells, to explore how snail-nematodes interactions may vary in response to city life. We examined shells from the generalist snail *Cepaea nemoralis* sampled in three urban areas in Belgium for trapped nematodes, and attempted to link this to urbanization and shell phenotypic traits. We found that even a small degree of urbanization led to large decreases in the rates of shell encapsulation, and that larger snails were more likely to contain trapped nematodes. However, we found no evidence that shell colour, which had been previously linked to immune function, was correlated to encapsulation rates. We discuss how between-population variation in encapsulation rates can result from urbanization-induced changes on the nematodes side, the snail side, or both, and suggest potential tests for future studies aiming to disentangle these mechanisms.

## Introduction

Urbanization is a major and all-encompassing human-induced environmental change, leading to changes in land use, local climate, soil imperviousness, light and chemical pollution… (Parris, 2016). The (often negative) impacts of these multivariate changes on biodiversity are increasingly well-documented: many species decline in cities, while some become successful “urban adapters”, leading to major restructuring of biological communities along urbanization gradients (e.g. McKinney, 2008; Piano et al., 2020; Fenoglio et al., 2020; Liang et al., 2023). In parallel, urbanization can also cause within-species phenotypic and genetic changes (Alberti et al., 2017; Szulkin et al., 2020; Diamond & Martin, 2021). Urbanization may also have second-order impacts by reshaping ecological interactions, if tightly connected species respond to environmental change in different ways (Theodorou, 2022). Such urbanization-induced changes in ecological interactions, in both positive and negative directions, have been recorded for plant-pollinator interactions (Liang et al., 2023), plant-herbivore and prey-predator interactions (Eötvös et al., 2018; Valdés-Correcher et al., 2022; Gámez et al., 2022; Korányi et al., 2022), as well as host-parasite interactions (Murray et al., 2019; Korányi et al., 2022).

Terrestrial molluscs (snails and slugs) are potentially valuable models in urban ecology and evolution, in part because of their limited movement abilities, which means they often cannot move to escape environmental changes. Like in many other taxa, urbanization can reshape molluscan communities (Lososová et al., 2011; Horsák et al., 2013; Barbato et al., 2017; Hodges & McKinney, 2018), and drive evolutionary responses in urban populations (Kerstes et al., 2019). Interestingly, in a comparative cross-taxon study of urbanization impacts, snail species richness were less negatively affected, compared to other more mobile groups (Piano et al., 2020). Land molluscs are hosts to a diverse array of metazoan parasites, including nematodes, flies, mites or trematodes (Barker, 2004; Segade et al., 2013; Żbikowska et al., 2020). How urbanization reshapes these interactions remains understudied, despite some of these parasites being of increasing veterinary interest (Aziz et al., 2016; Giannelli et al., 2016).

Snails and slugs can defend themselves against metazoan parasites through a variety of behavioural (Wilson et al., 1999; Wynne et al., 2016; Rae, 2023) or immune responses (Furuta & Yamaguchi, 2001; Scheil et al., 2014; Coaglio et al., 2018). Among the latter, it has been discovered that land molluscs can use their shells to trap parasitic nematodes, killing them and fusing them to the inner shell surface (Rae et al., 2008; Williams & Rae, 2015; Rae, 2017). This ability seems phylogenetically widespread, even present in slugs with vestigial shells (Rae et al., 2008; Rae, 2017), and could therefore provide a relatively easy to access record of ecological interactions. Following anecdotal records of mites and trematodes encapsulated in shells, it has further been suggested that this shell encapsulation might extend to other metazoan parasites (Dahirel et al., 2022; Gérard et al., 2023). However, given how rare these non-nematode records are, they may be merely by-products of a defence mechanism targeted towards nematodes, rather than evidence of a more generalized defence response (Gérard et al., 2023). The few snail species in which this phenomenon has been studied across multiple populations show that the prevalence of individuals trapping nematodes can vary widely between sites (Rae, 2017; Rae, 2018; Cowlishaw et al., 2019), but there has been no attempt, to our knowledge, to assess whether this variation could be non-random with respect to environmental context.

To that end, we combine here publicly available and standardized urbanization metrics with observations of field-collected snails across three cities in Belgium, using the grove snail *Cepaea nemoralis* (Gastropoda, family Helicidae) as a model. Like other helicids, *C. nemoralis* can encapsulate and trap parasitic nematodes in its shell (Williams & Rae, 2016; Rae, 2017; Dahirel et al., 2022; Gérard et al., 2023). This snail is also common both outside and within cities (Kerstes et al., 2019), and therefore a very suitable model to study variation in encapsulation rates, whether it is due to urbanization or to spatial (between-cities) differences. Furthermore, the shell colour variation that made *Cepaea* species iconic models in evolutionary biology (Jones et al., 1977; Ożgo, 2009) may also influence their immune response, with some evidence that darker morphs mount better defences against nematodes (Dahirel et al., 2022; but see Scheil et al., 2014). On the other hand, this morph variation in resistance might not translate to shell encapsulation (Williams & Rae, 2016; Dahirel et al., 2022). However, existing comparisons were either limited to one type of colour variation (banding pattern only, Dahirel et al., 2022), or analysed experimental infections by one model nematode (Williams & Rae, 2016); we here test whether this remains true when analysing naturally occurring snail-nematode interactions and accounting for more dimensions of shell colour variation.

## Methods

### Site selection and sampling

We searched for *Cepaea nemoralis* snails from early October to mid-November 2022 in and around the urban areas of Brussels, Ghent and Leuven in Belgium (**Fig. 1**). Potential sites were selected based on pre-existing online crowdsourced records (iNaturalist contributors & iNaturalist, 2024) combined with personal observations and virtual fieldwork using Google StreetView to identify suitable habitats (based on Falkner et al., 2001). We visited 36 sites chosen to be roughly balanced between the three cities (including their surrounding areas; Brussels: 13 sites, Ghent: 13 sites, Leuven: 10 sites). In each site, we sampled living snails by hand during visual search, in a radius of up to 50 m around a designated site centroid (though search was *de facto* mostly concentrated within a 20 m radius). Individuals were mainly searched in known favourable micro-habitats, i.e. on tall herbs and shrubs, under piled wood and cardboard or loose rocks, or on fences, walls, and tree trunks (Falkner et al., 2001). Field identification of *Cepaea nemoralis* snails is easy based on shell shape, size and colour (Cameron, 2008). We only collected adults, which can easily be separated from subadults by the presence of a reflected shell lip marking the end of shell growth (Cameron, 2008). Each site was visited by 1 to 3 people (mean: 2.03) for a duration of 5 to 30 person-minutes (mean: 15). We collected a total of 298 snails from 28 of the 36 sites visited (Brussels: 9 sites, Ghent: 10 sites, Leuven: 9 sites). However, 2 shells were lost before examination for parasites due to handling errors, and another shell was accidentally broken for parasite examination before photographs or size measurements could be done; this led to a final complete dataset of 295 snails in 28 sites. For each of these 28 sites, the nearest neighbouring site with snails found was between 153 and 1516 m away (mean: 768 m), which is in any case farther than the maximal dispersal distances (Kramarenko, 2014), indicating that even nearby sites could be considered separate populations.

**Figure 1.**
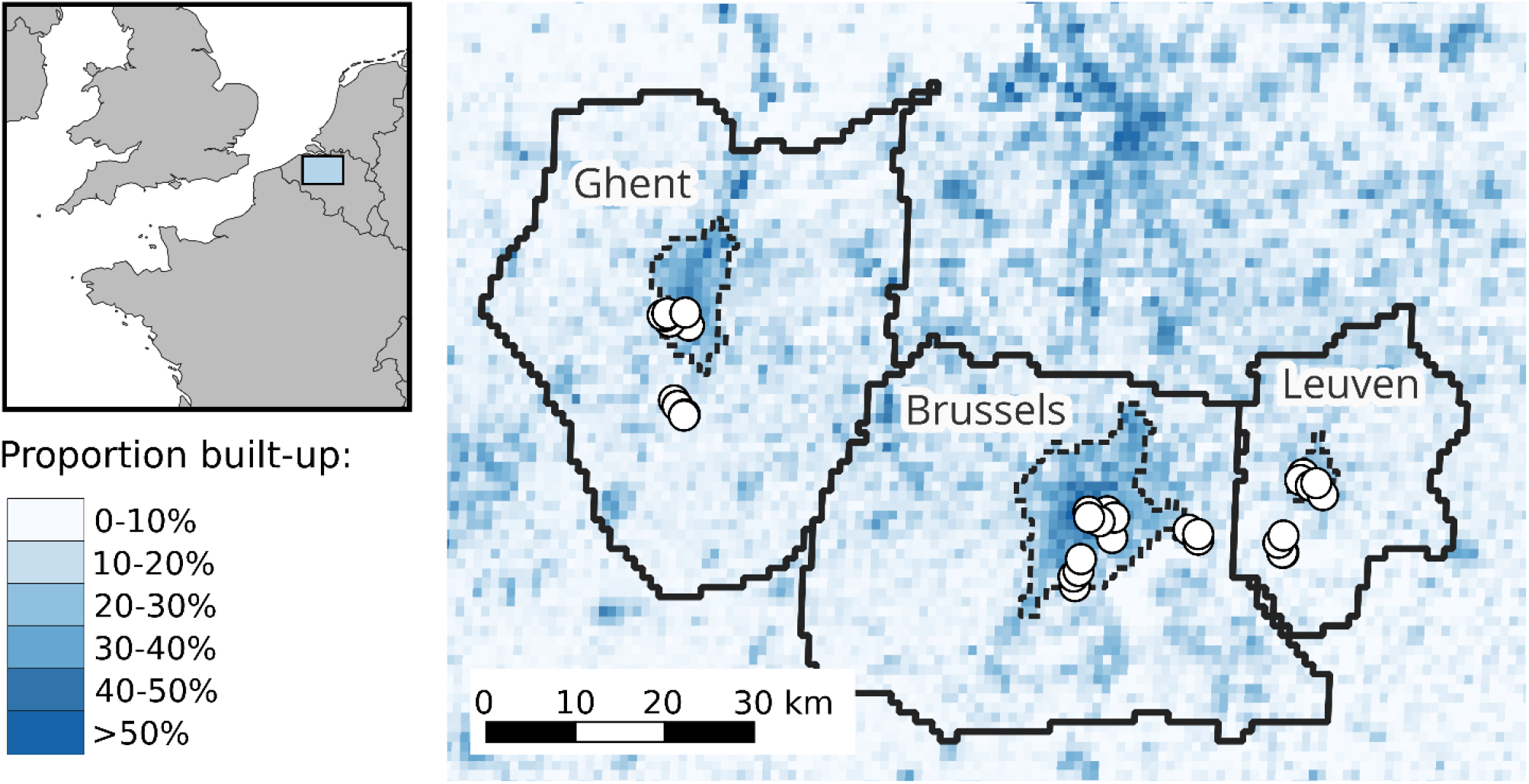
Location of study sites within western Europe and central Belgium. The Functional Urban Areas (roughly corresponding to commuter zones, Schiavina et al., 2019; Moreno-Monroy et al., 2021) that were used to link each site to a city are also displayed as solid black lines, while the corresponding core urban areas (Urban Centres *sensu* Eurostat (European Commission), 2021; Schiavina, Melchiorri, et al., 2023) are displayed with dashed lines.

### Urbanization metrics

It is well-known that urban environmental changes are complex and multivariate (e.g. Parris, 2016); however, given our relatively low number of sites, and the risk of collinearity between urban metrics, we decided to use simple overarching metrics focused on building presence and human population density. We assessed urbanization at each site using raster layers from the Global Human Settlement Layer project for the year 2020 (https://ghsl.jrc.ec.europa.eu/, Joint Research Centre (European Commission), 2023). We first used built-up surface (GHS-BUILT-S) and population density (GHS-POP) at 100 m and 1000 m resolutions (Pesaresi & Politis, 2023; Schiavina, Freire, et al., 2023). The former spatial scale matches the scale of maximal dispersal movements over timespans of up to a couple years in helicid snails, while the latter is closer to the scale of longer term (over several decades) population spread (Kramarenko, 2014). As an additional categorical metric, we also used the Degree of Urbanization as recorded in the Settlement Model layer (GHS-SMOD, available only at 1000 m resolution, Eurostat (European Commission), 2021; Schiavina, Melchiorri, et al., 2023). At the highest level of classification, the standardized Degree of Urbanization methodology mainly uses population density and contiguity rules to classify grid cells as either part of a continuous high-density Urban Centre, as low-density rural cells or as intermediate peri-urban/suburban cells. For each site and urbanization metric, we recorded the value of the corresponding grid cell. Interestingly, Degree of Urbanization classes, primarily based on population density, divide our sites in almost the same non-linear way as another, independent, three-level classification based on built-up surfaces used in previous urban ecology studies in the study region (e.g. Piano et al., 2020) (**Supplementary Material S1**).

### Snail shell analysis

Snail size was measured using a caliper as the shell greatest diameter (to the nearest 0.1 mm). Snail shell colour morphs were scored following e.g. Cain (1988) for background colour (from lighter to darker: yellow, pink or brown), number of dark bands (0 to 5 bands) and on the presence or absence of band fusions (which increase the proportion of the shell covered by dark bands). Snails were killed by first inducing dormancy at 6°C, then by freezing at -20°C. We removed bodies from shells with forceps and lightly cleaned shells with water (bodies were stored in ethanol for separately planned studies). We then broke each shell into fragments using forceps, examined fragments under a binocular microscope, and recorded all animals found encapsulated within the shell as in e.g. Gérard et al. (2023). A total of 606 nematodes were found in 104 shells (**Fig. 2**); we found no mites, trematodes or other parasites in any of the shells. Shells with nematodes contained 5.83 nematodes on average (SD: 9.95, range: 1-58). As this method is destructive, we took standardised photographs of the shells beforehand (dorsal and apertural views following Callomon, 2019) for archival and potential future studies.

**Figure 2.**
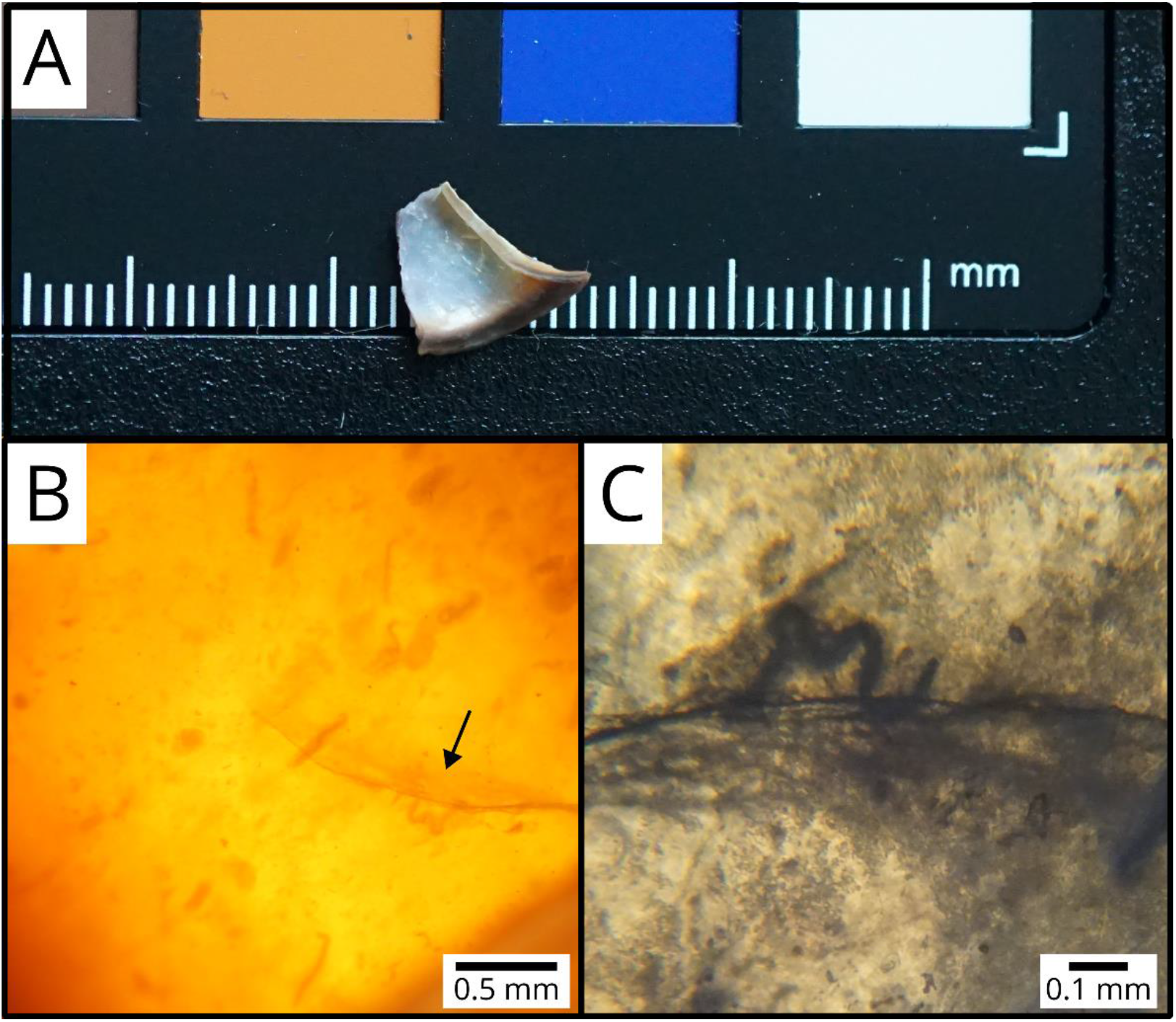
Fragment of a *Cepaea nemoralis* shell (A) containing encapsulated nematodes (B, C). The arrow in (B) points to the nematode shown in (C).

### Statistical analysis

All analyses were done in R version 4.3.2 (R Core Team, 2023), with the help of the *tidyverse* (Wickham et al., 2019) and *sf* (Pebesma, 2018) packages for data processing, as well as additional packages detailed below for model fitting and exploration.

We analysed the probability a shell contained nematodes as a binary yes/no response at the individual level, using Generalized Linear Mixed Models (GLMMs) (binomial family, logit link). We ran six models; the first five all included shell size, shell morph traits (background colour, band number and fusion), urbanization and city identity (Brussels, Ghent or Leuven) as fixed effects, only differing by which urbanization metric they used (among the five described above in **Urbanization metrics**). Given our sample size, we did not include interactions between our explanatory variables, especially as we had no *a priori* hypotheses regarding these (but see **Discussion**). Numeric predictors were centred and scaled to unit 1 SD. Sampling site was included as a random intercept. The sixth model was a “null” model, identical to the other ones except that it did not include an urbanization metric. We ran our models using the *glmmTMB* package (Brooks et al., 2017), and then used AICc to compare them. As one model largely outperformed the others (see **Results**), we did all further analyses on that best model.

We checked for residual spatial autocorrelation using a spline correlogram (*ncf* package, Bjornstad, 2022), and found no evidence of spatial structure in the best model. We then used the *car* (Fox & Weisberg, 2019) and *emmeans* (Lenth, 2023) packages to test for overall effects of our variables in the best model and to run (Tukey-corrected) pairwise comparisons, respectively. Finally, we estimated the marginal and conditional *R*^2^ (Nakagawa & Schielzeth, 2013) as measures of the proportion of variation explained by fixed effects 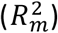 and both fixed and random effects 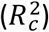 respectively (using the delta method, Nakagawa et al., 2017).

## Results

The model using the categorical Degree of Urbanization (GHS-SMOD) as an urbanization metric outperformed all other models based on AICc (**Table 1**). Fixed effects and random effects explained similar amounts of variance 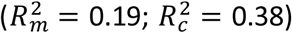. The probability that a shell had trapped nematodes was dependent on urbanization level (*χ*^2^ = 15.97, df = 2, *p* = 3.40 × 10^−4^) but did not vary significantly between cities (*χ*^2^ = 3.89, df = 2, *p* = 0.14). Snails from rural sites were more likely to contain nematodes than snails from intermediate and Urban Centre populations (**Fig. 3**; rural - intermediate difference on the logit scale ±SE: 3.71 ±0.95 ; rural - Urban Centre difference: 2.34 ±0.77). Larger shells were more likely to contain nematodes (*χ*^2^ = 4.17, df = 1, *p* = 0.04 ; standardised coefficient *β* = 0.35 ±0.17). There was no clear evidence that any of the shell colour traits affected encapsulation rates (background colour: *χ*^2^ = 2.17, df = 2, *p* = 0.34; band number: *χ*^2^ = 1.90, df = 1, *p* = 0.17 ; fusion: *χ*^2^ = 0.17, df = 1, *p* = 0.68).

**Table 1.**
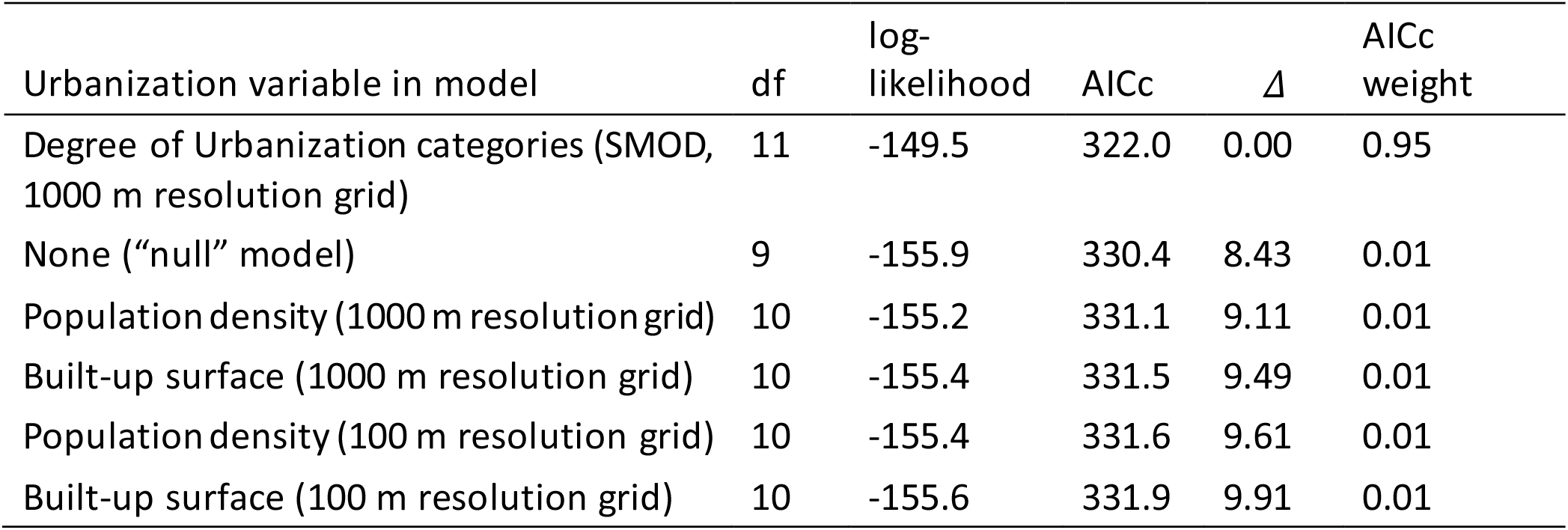
Model selection table for the effect of urbanization on shell encapsulation rates. All models otherwise include effects of city identity, shell size and shell morph (background colour, number of bands and band fusion).

**Figure 3.**
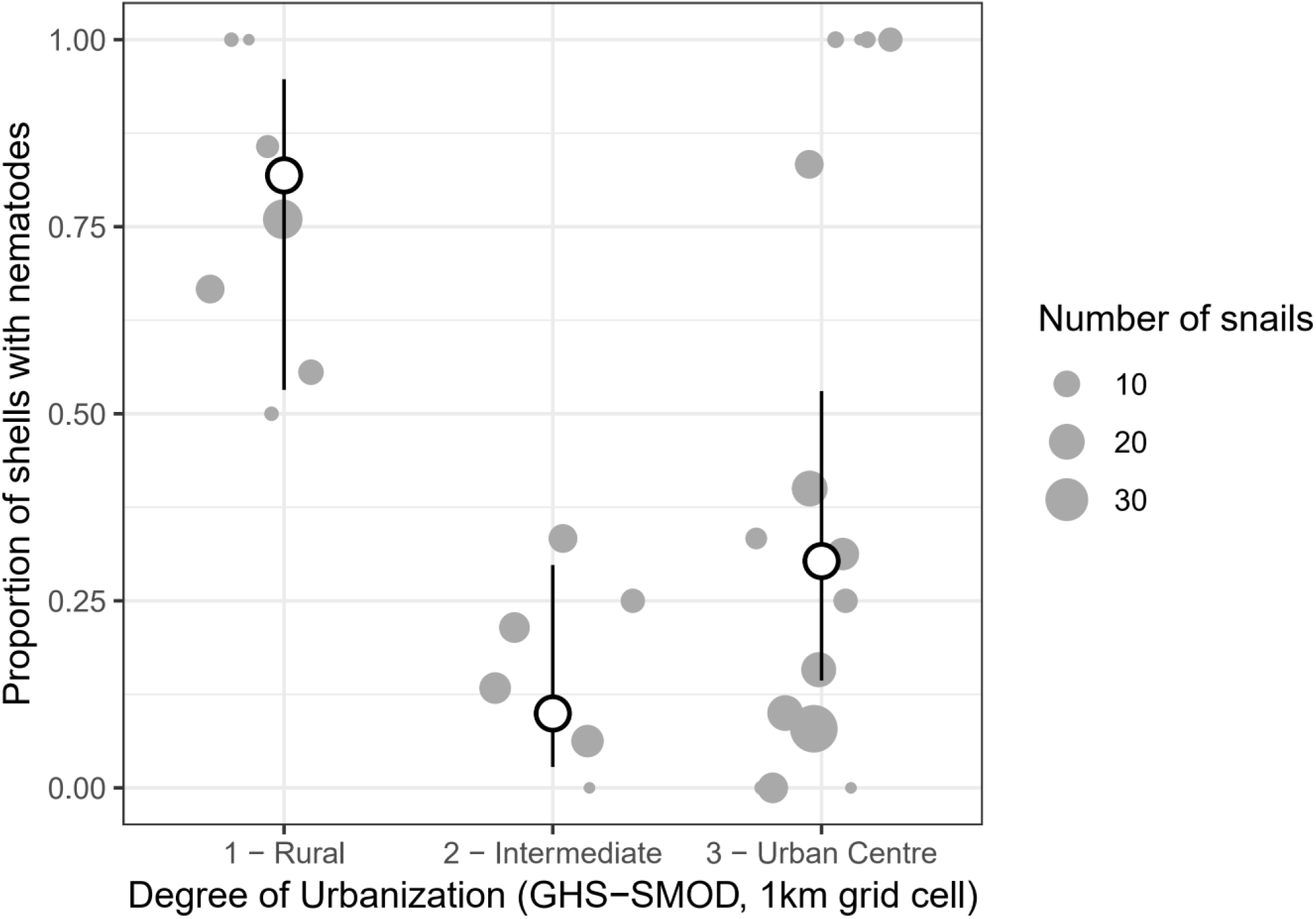
Effect of the Degree of Urbanization in 1000 m grid cells on the probability a snail shell contained encapsulated nematodes. Grey dots are observed proportions per population, with the size of the dot proportional to the number of snails; white dots (and error bars) are estimated marginal means from the best model (and their 95% confidence intervals), with the effects of the other predictors averaged out.

## Discussion

We found that the prevalence of *Cepaea nemoralis* snails encapsulating nematodes in their shell as a defence mechanism were partly driven by environmental conditions, with lower trapping rates in more urban sites (**Fig. 3**). This effect was better explained by a categorical classification of the Degree of Urbanization, rather than by linear effects of continuous urbanization variables. This indicates that the response to urbanization is non-linear, as the increases in population density/built-up rates needed to go from a rural to an intermediate area are much smaller than those needed to go from intermediate to Urban Centre, and most of the variation in density/built-up is within Urban Centres rather than between categories (**Supplementary Material S1**).

A difficulty for interpreting our results is that nematodes trapped in shells accumulate with time (Williams & Rae, 2015; Rae, 2017), meaning that as they may have endured more infections, older snails may be more likely to have them. If the urban heat island influences snail survival rates (Wolda, 1967, Manoli et al., 2019), then our urban-rural differences in nematodes trapped might merely reflect differences in average snail age/survival. Age estimation in terrestrial snails is challenging (Pollard et al., 1977; Williamson, 1979), and any age proxy is likely to be influenced by local conditions, making it useless to compare age between populations without thorough calibration studies. However, while there is substantial population variation, the number of nematodes found in infected shells does not decrease in more urbanized sites (**Supplementary Material S2**), contrary to what we would expect if variation in shell encapsulation was primarily explained by variation in time available to accumulate nematodes.

If we assume that our results reflect differences in snail-nematodes interactions between urban and non-urban areas, several mutually non-exclusive mechanisms may explain why urban *Cepaea nemoralis* shells are less likely to trap nematodes. Each of these mechanisms directly suggests potential tests for future studies:

- First, snail parasitic nematodes infecting *C. nemoralis* may be less abundant in cities. Many nematodes known to infect land snails have at least one free-living life stage in the soil, and some are facultative parasites (Morand et al., 2004; Pieterse et al., 2017). Increasingly impervious substrates in cities (Parris, 2016) may deprive these of habitat critical for their life cycle. Where habitat is available, soil nematode communities are profoundly altered by urbanization, like other taxa (Li et al., 2022; Gong et al., 2024). However, this does not lead to overarching declines in nematode abundance; rather, some trophic groups decline while others thrive (Li et al., 2022; Gong et al., 2024). Unfortunately, detailed information on nematodes parasitizing animals is typically lacking from these analyses; soil sampling specifically targeting parasitic nematodes (Jaffuel et al., 2019) would be here particularly useful. The few studies available are mixed on the effects of urbanization on the infection of land molluscs by parasitic nematodes. In Wales, urban and suburban slugs are more, not less, likely to be infected by *Angiostrongylus vasorum* compared to rural ones (Aziz et al., 2016). By contrast, data from Andrus et al. (2022) spanning urban and non-urban sites suggest that the prevalence of nematode infection may be slightly lower in urban molluscs, although they did not themselves analyze the effect of urbanization. Both studies however analyzed nematode prevalence in molluscs, not their abundance/availability in the urban environment.
- Second, individual differences in behaviour, especially space-related behaviour, may lead to differences in the risk of encountering and then being infected by parasites (Barber & Dingemanse, 2010). Habitat loss and fragmentation associated with urbanization are expected to exert strong selection pressures on movement and space use (Cote et al., 2017). If this results in lower movement in urban snail populations, this might then reduce their encounter rates with parasites. In the snail *Cornu aspersum*, urbanization does not lead to reduced habitat boundary-crossing behaviour (Dahirel et al., 2016), although that is only one component of mobility. Urbanization-induced increases in temperature may also alter the frequency at which snails hide into shelters or climb above the substrate (Rosin et al., 2018), and potentially again the risk of encountering parasites. The picture is complicated by behaviour-parasite feedbacks, where while host behaviour shapes infection risk, infection can then alter host behaviour in turn (Ezenwa et al., 2016). In *Cepaea nemoralis*, nematode infection itself might lead to reduced movement propensity, but only in some morphs (Dahirel et al., 2022). More studies of movement behaviour across urbanization gradients are here needed.
- Third, shell encapsulation rates are not direct records of snail-nematode interactions, but rather informative on the host’s ability to mount a defence in such interactions. This defence is not always effective, as field-caught snails sometimes show active infections but zero shell-trapped nematodes (see e.g. data in Dahirel et al., 2022). If immune response declines with urbanization, then this alone could explain our results even in the absence of changes in nematode communities. In vertebrates, urban living can lead to both depressed or stimulated immune function, depending on taxon and context, especially food availability (Murray et al., 2019; Minias, 2023). In terrestrial molluscs, chemical pollutants seem to negatively impact many, but not all, physiological components of immune defence (Radwan et al., 2020). The exact physiological pathways involved in shell encapsuIation in land molluscs remain however unstudied, to the best of our knowledge.

Interestingly, nematode encapsulation prevalence was seemingly more variable between Urban Centre populations than between populations in the other urban categories, with a few sites having observed prevalences largely above the predicted mean (**Fig. 3**). While this may simply be sampling variability as these populations have very small sample sizes, this suggests that there could be non-random within-city variability in snail-nematode interactions. As a first post-hoc exploration, we have re-run the models with continuous urbanization variables as predictors, using only the Urban Centre subset of sites (see **Data and code availability**). After accounting for phenotype and city of origin, we found no indication that built-up or population density levels influenced prevalence within the Urban Centre category. Nonetheless, cities remain highly heterogeneous environments even beyond built-up and population density; for instance, within-city variation in vegetation, mediated in part by neighborhood-level socio-economic differences, may shape biodiversity, including species interactions (e.g. Martin et al., 2024). While we are not able to identify the causes of this heterogeneity in our current dataset, as we are hampered by our small number of sites per Urban Centre, future studies designed to target this within-city heterogeneity may uncover more on the fine-scale drivers of snail responses to parasites.

On the individual phenotype side, larger shells were more likely to contain trapped nematodes. If shell size also varied in response to urbanization, then this could open an indirect pathway linking urbanization to encapsulation mediated by snail size, potentially accentuating or dampening the direct effect we describe above. However, we found no clear effect of urbanization on *C. nemoralis* shell size (**Supplementary Material S3**). In addition and as a post-hoc exploration, we re-ran our model set adding size × urbanization interactions, and found no significant interaction, and no evidence that the urbanization effect changed in response (**Supplementary Material S4**). The relationship between size and nematode encapsulation could be the result of survivor bias alone, if larger snails are more likely to survive infection. However, and although we cannot exclude that other nematodes have larger effects, experimental nematode infections by *Phasmarhabditis* are almost never lethal in adult *Cepaea nemoralis*, contrary to other snail species (Wilson et al., 2000; Williams & Rae, 2016). Other potential explanations for this result can be sorted along three non-exclusive lines, similar to the mechanisms suggested above to explain the effect of urbanization:

- Larger snails might harbour larger parasite infections (e.g. Daniels et al., 2013), which would increase the likelihood that some nematodes are trapped. However, there is no link between nematode abundance in active infections and snail size in *C. nemoralis* (Dahirel et al., 2022), and no clear effect of shell size on the number of nematodes trapped in the present study (**Supplementary Material S2**).
- If large and small snails differ in their space use, they might also differ in their parasite exposure risk. Evidence for a link between shell size and space use is mixed in *Cepaea nemoralis*, and this may depend on the scale of the movements in question (short-term routine vs. dispersal movements; Oosterhoff, 1977; Dahirel et al., 2022).
- Finally, small and large snails may differ in their immune defence abilities. Comparative studies suggest that large and small snail species and subspecies differ in their immune strategies at the physiological level (Russo & Madec, 2011, 2013). However, the range of body size and life history variation is much larger in these scenarios than among adults of *C. nemoralis*, limiting the transferability of these results. More physiological studies focused on within-, rather than among-species variation may help understand better this link between body size and encapsulation rates.

In contrast to shell size, we found no relationship between any of the shell colour traits and nematode trapping rate. This confirms experimental results from Williams & Rae (2016) using infections by *Phasmarhabditis hermaphrodita*. However, colour morphs do differ in active infection rates or other aspects of immune response in *C. nemoralis* (Dahirel et al., 2022) and other polymorphic snails (Scheil et al., 2013, 2014). This discrepancy may indicate that shell encapsulation is driven by different physiological pathways than other components of snail immune defence.

Beyond the effects of phenotype or environment, whether and how the prevalence of nematodes trapped in shells is correlated with rates of active parasite infections remains an open and complex question (which we could not tackle here as snail bodies were reserved for other investigations). If variation in snail-nematode interactions is driven by e.g. variation in nematode density in the environment, we may expect a positive correlation, as higher nematode densities should drive up rates of both shell encapsulation (Rae, 2018) and active infection (although if encapsulation is highly effective, it may end up suppressing dose-dependent effects on active infection, Williams & Rae, 2015). On the other hand, if variation is mostly driven by snail immune response, we may expect a negative correlation: snails with more effective immune systems may be more likely to successfully trap nematodes in shells while being less likely to harbour active infections. While this would need to be validated, the strength and direction of between- and within-sites correlations between active infections and shell-trapped nematodes may provide useful indicators of the main drivers of snail-nematodes interactions in response to city life.

We acknowledge that the relatively small size of our sample does not allow us to draw firm causal conclusions. Nonetheless, we hope our results may encourage larger studies regarding host-parasite interactions in land molluscs in the context of environmental change. As new technical developments such as micro-CT imaging allow non-destructive analyses of snail shells (Falkingham & Rae, 2021), these may extend to using museum and other natural history collections to understand how interactions vary in space and time (Cowlishaw et al., 2019), reaffirming their value for urban ecology and evolution (Shultz et al., 2020).

## Author contributions

Initial study idea: MD, DB. Site selection and fieldwork: MD, HR, KDW. Shell data collection: HR, after initial training by MD. Data analysis: MD, after preliminary analyses by HR. Initial manuscript draft: MD. All authors contributed critically to edits and gave final approval for publication.

## Data and code availability

Data and R scripts to reproduce all analyses presented in this article, as well as a copy of the Supplementary Materials, are available on Github (https://github.com/mdahirel/HELICITY-2022_shell-nematodes) and archived in Zenodo (DOI: https://doi.org/10.5281/zenodo.10794928).

