## Supplementary Material for "Urban *Cepaea nemoralis* snails are less likely to have nematodes trapped within their shells"

### *Cepaea nemoralis* snails are less likely to trap nematodes within their shells in urban areas - supplementary material

Maxime Dahirel, Hannah Reyné, Katrien De Wolf, Dries Bonte

#### S1 - Variation in human population density and in built-up surfaces between Degrees of Urbanization

We show here (**Figs S1-1 and S1-2**) the relationships in our sites between the Degree of Urbanization classes (Eurostat (European Commission), 2021) and the corresponding values of human population density and built-up cover, both from 1000 m resolution rasters.

- For both population density and built-up, the differences between the classes scale non-linearly, with the difference between the average Urban Centre site and the average "intermediate" site being larger than the difference between the average intermediate and rural sites;
- Urban Centre sites contain a much broader range of variation in either variable, compared to intermediate and rural sites;
- Interestingly, the differences in built-up surfaces between the three Degree of Urbanization classes map almost exactly to built-up thresholds designed and used independently in a series of urban ecology studies from Belgian sites (e.g. Merckx et al., 2018; Piano et al., 2020) (**Fig. S1-2**). Degree of Urbanization classes can therefore (*for our data*) be interpreted as encoding both variation in density and in built-up surfaces.

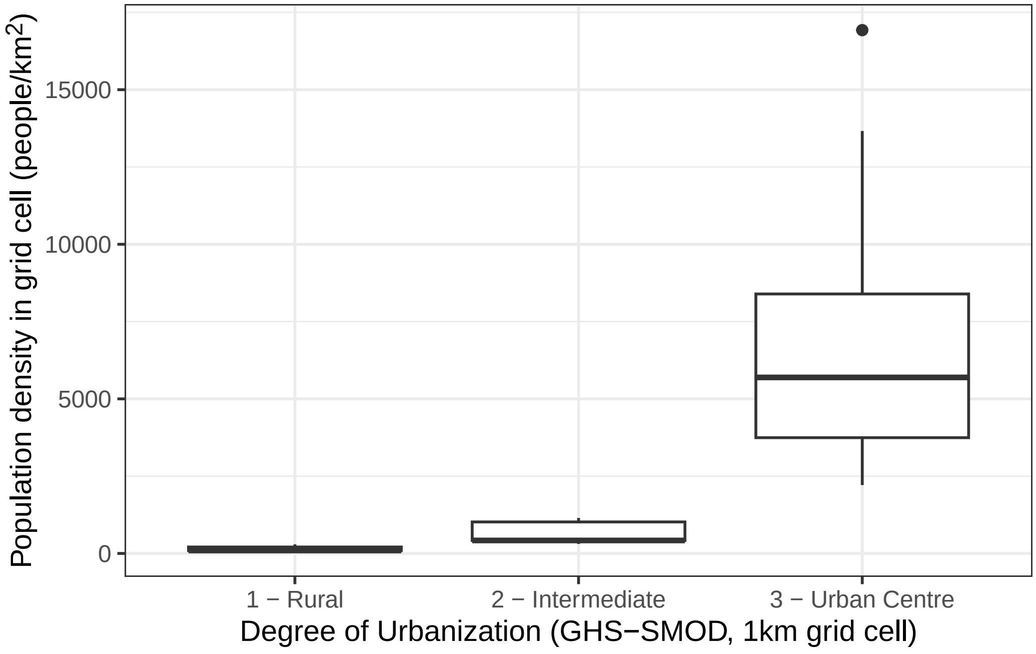


**Figure S1-1.** Relationship between Degree of Urbanization classes and human population densities, for all 36 visited sites (including sites with no snails found).

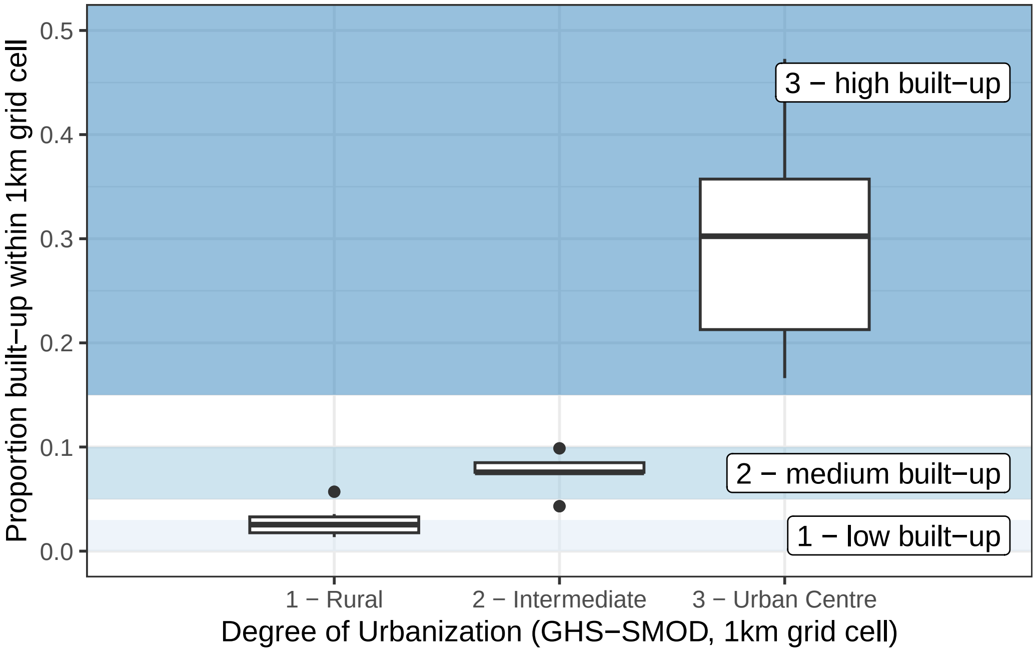


**Figure S1-2.** Relationship between Degree of Urbanization classes and built-up surfaces, for all 36 visited sites (including sites with no snails found). Shaded areas and corresponding labels mark the limits of the three urban classes used in studies from the Belgian SPEEDY project, including for instance Merckx et al. (2018) or Piano et al. (2020).

#### S2 - Intensity of nematodes trapped in shells

Shells that do contain nematodes may also vary in the numbers of nematodes they harbour (in the context of active infections, this is termed "intensity," Bush et al., 1997). However, given the relatively low number of shells with nematodes (104) especially relative to the number of sites, and given the expected high dispersion of these numbers (e.g. Rae, 2017), we do not consider our intensity dataset to be sufficiently large to be given consideration in the main text. The partial analysis of these data we present below is therefore intended for indicative purposes only.

Using the subset of shells where nematodes were detected, we ran a negative binomial GLMM on the number of nematodes per shell - 1, with the same fixed and random effect structure as the best model for nematode prevalence (see **main text**). We find no clear indication that these numbers differed between urbanization levels (**Table S2-1**, **Fig. S2-1**), or in relation to shell phenotype (**Table S2-1**). The conditional $R^2$ (proportion of variance explained by both fixed and random effects) is 0.42 while the marginal $R^2$ (fixed effects only) is 0.06, indicating that there are substantial population differences in nematode numbers, even though they are not explained by urbanization, city of origin or shell phenotype. Indeed, the mean number of nematodes found in shells varied substantially between populations, ranging from 1 to 22.5 nematodes.

**Table S2-1.** Analysis of Deviance for the effect of urbanization and shell phenotype on the number of nematodes per shell (on the subset of shells containing at least one nematode).

| variable | $\chi^2$ | df | $p$ |
|---|---|---|---|
| Degree of Urbanization | 0.02 | 2 | 0.99 |
| City | 0.48 | 2 | 0.79 |
| Shell background colour | 0.18 | 2 | 0.91 |
| Number of bands | 0.76 | 1 | 0.38 |
| Band fusion | 0.82 | 1 | 0.36 |
| Shell size | 1.89 | 1 | 0.17 |

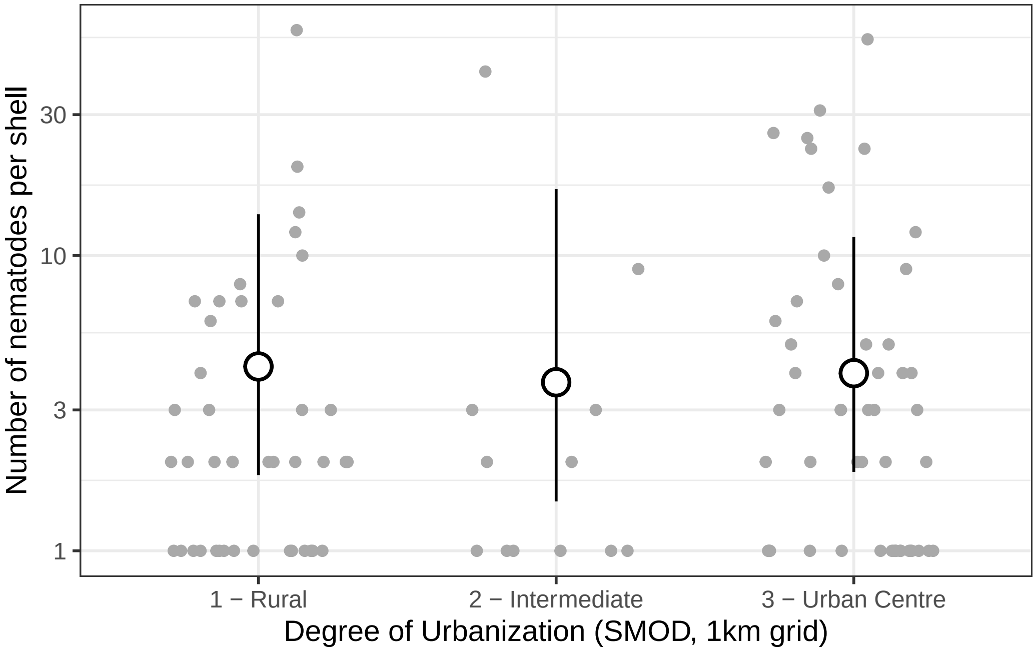


**Figure S2-1.** Relationship between the Degree of Urbanization in 1000m grid cells and the number of nematodes *in shells that contain at least one nematode*. Grey dots are observed values; white dots (and error bars) are estimated marginal means from the best model (and their 95% confidence intervals), with the effects of city identity and phenotypic traits averaged out. Note the $\log_{10}$ scale on the $y$ axis.

#### S3 - Potential relationship between urbanization and shell size

In the main analyses, we showed that both urbanization and snail shell size have an effect on the probability of having encapsulated nematodes. If snail size were also influenced by urbanization (as in e.g. Theodorou et al., 2021), this would open the way for both direct and indirect (mediated by size) effects of urbanization on nematode encapsulation. In that case, we would need to combine both to have an idea of the total effect of urbanization on nematodes.

To check this, we ran a linear mixed model to test for a relationship between urbanization levels (Degree of Urbanization categories) and body size. The model included city of origin as a covariate, and population of origin as a random intercept.

We found no evidence of an effect of urbanization on shell size (**Fig. S3-1**, **Table S3-1**). However, snails from Leuven were consistently smaller than snails coming from the other two cities, irrespective of urbanization level (**Fig. S3-2**, **Table S3-1**).

**Table S3-1.** Analysis of Deviance for the effect of urbanization on shell size (all *Cepaea nemoralis* independently of nematode encapsulation).

| variable | $\chi^2$ | df | *p* |
|---|---|---|---|
| Degree of Urbanization | 2.06 | 2 | 0.36 |
| City | 10.42 | 2 | 0.01 |

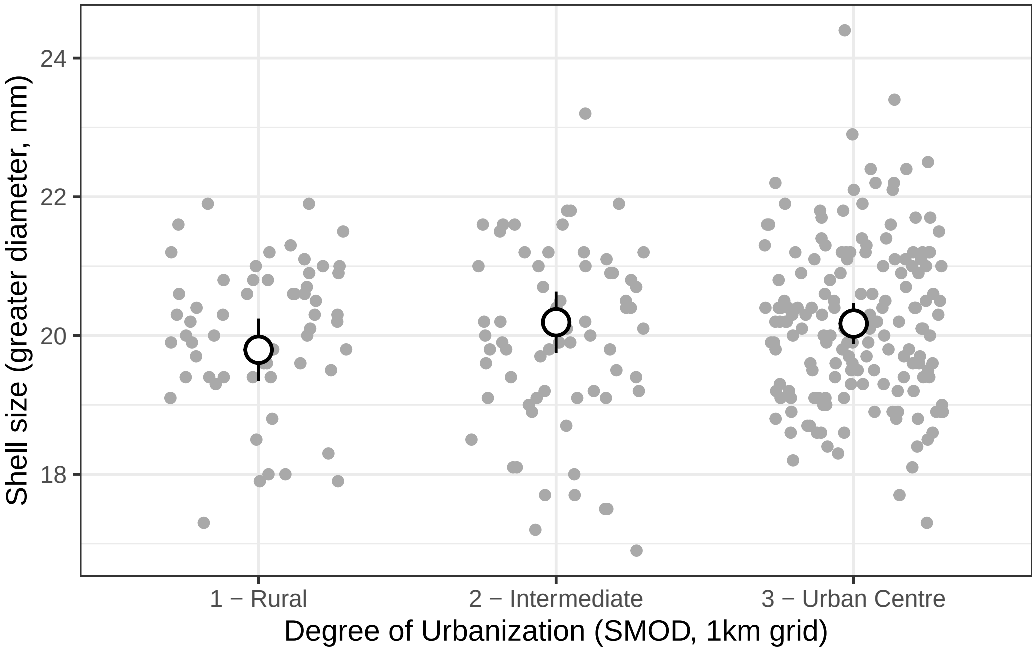


**Figure S3-1.** Relationship between the Degree of Urbanization in 1000m grid cells and snail shell size. Grey dots are observed values; white dots (and error bars) are estimated marginal means from the best model (and their 95% confidence intervals), with the effects of city identity averaged out.

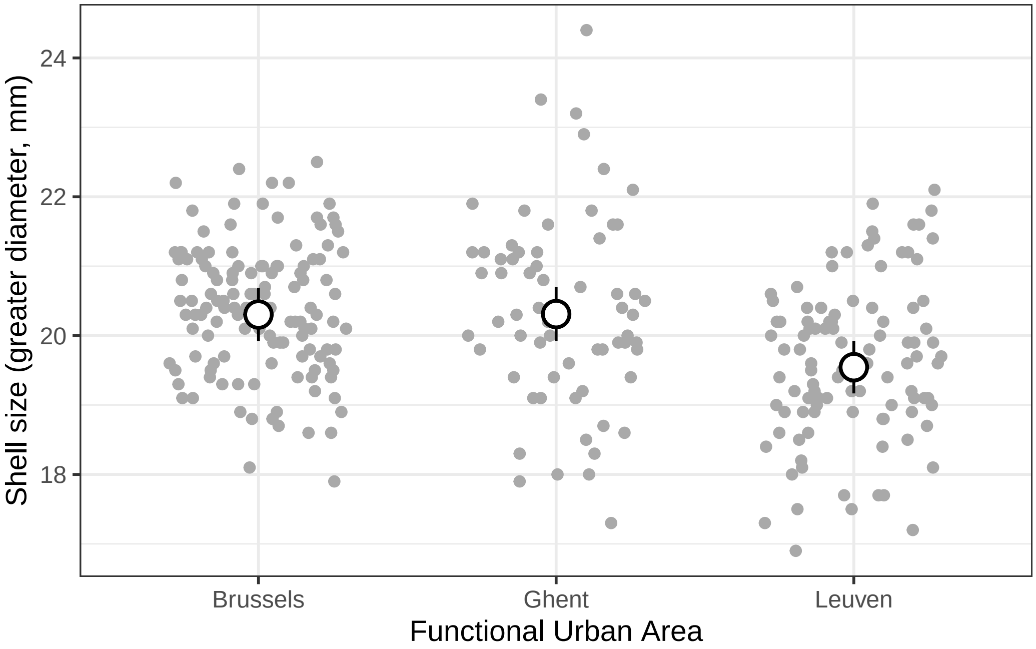


**Figure S3-2.** (left) Relationship between city (Functional Urban Area) of origin and snail shell size. Grey dots are observed values; white dots (and error bars) are estimated marginal means from the best model (and their 95% confidence intervals), with the effects of urbanization levels averaged out.

#### S4 - Potential interactions between urbanization and shell size

Another way for urbanization to have complex indirect effects through size is that if the effect of shell size on nematode prevalence depends on urbanization, i.e. if there are interactions between the two.

In this post-hoc analysis, we explored this possibility by re-running our model set, this time adding size × urbanization interactions to all models (except of course the "null" model with no urbanization effect).

The model using the categorical Degree of Urbanization (GHS-SMOD) again outperformed all other models based on AICc (**Table S4-1**). In this model, fixed effects and random effects again accounted for similar amounts of variance ($R^2_m = 0.23$; $R^2_c = 0.40$).

There was no significant size × urbanization interaction ($\chi^2 = 3.06$, df = 2, $p = 0.22$).

The probability that a shell contained trapped nematodes was still dependent on urbanization level ($\chi^2 = 16.76$, df = 2, $p = 2.30 \times 10^{-4}$). The predicted values for each urbanization level were nearly identical to those obtained from the original model in the main text (**Fig. S4-1**;

rural - intermediate difference on the logit scale ± SE: 3.82 ± 0.95 ; rural - Urban Centre difference: 2.36 ± 0.76).

Importantly, with the addition of the (non-significant) size × urbanization interaction, the main effect of shell size itself became non-significant too ($\chi^2$ = 1.44, df = 1, $p$ = 0.23). The other effects that were initially non-significant remained so (city: $\chi^2$ = 4.11, df = 2, $p$ = 0.13; background colour: $\chi^2$ = 1.97, df = 2, $p$ = 0.37; band number: $\chi^2$ = 1.67, df = 1, $p$ = 0.20 ; fusion: $\chi^2$ = 0.21, df = 1, $p$ = 0.65).

**Table S4-1.** Model selection table for the effect of urbanization on shell encapsulation rates, for the modified model set with size × urbanization interactions added. As in the main text, all models otherwise include effects of city identity, shell size and shell morph (background colour, number of bands and band fusion).

| Urbanization variable in model | df | log-likelihood | AICc | Δ | AICc weight |
|---|---|---|---|---|---|
| Degree of Urbanization categories (SMOD, 1000 m resolution grid) | 13 | -147.9 | 323.1 | 0.00 | 0.90 |
| Population density (1000 m resolution grid) | 11 | -153.4 | 329.7 | 6.62 | 0.03 |
| None (“null” model) | 9 | -155.9 | 330.4 | 7.34 | 0.02 |
| Built-up surface (1000 m resolution grid) | 11 | -153.8 | 330.5 | 7.36 | 0.02 |
| Built-up surface (1000 m resolution grid) | 11 | -154.3 | 331.5 | 8.44 | 0.01 |
| Population density (100 m resolution grid) | 11 | -154.8 | 332.6 | 9.48 | 0.01 |

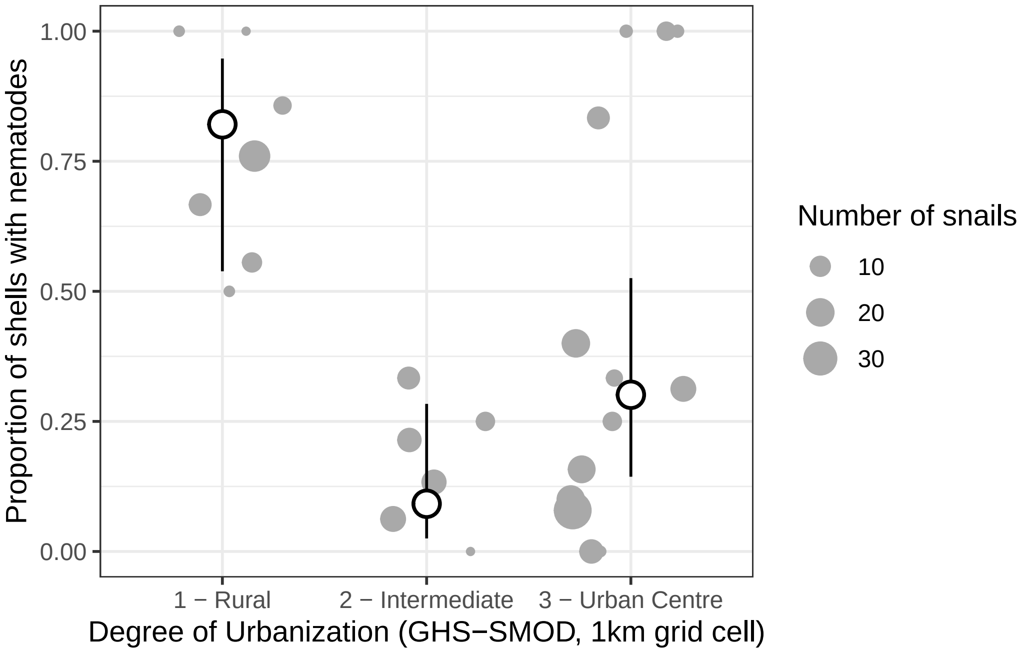

**Figure S4-1.** Effect of the Degree of Urbanization in 1000 m grid cells on the probability a snail shell contained encapsulated nematodes, based on the modified model with size × urbanization interactions added. Grey dots are observed proportions per population, with the size of the dot proportional to the number of snails; white dots (and error bars) are estimated marginal means from the best model (and their 95% confidence intervals), with the effects of the other predictors averaged out.
